# Weak Task Synchronization of Default Mode Network in Task Based Paradigms

**DOI:** 10.1101/2021.04.09.439206

**Authors:** Vaibhav Tripathi, Rahul Garg

## Abstract

Default mode network (DMN) are the areas of the brain that get deactivated upon the presentation of externally focused, attention demanding tasks. Interestingly, these areas are consistently deactivated irrespective of the nature of the task. Recent evidence shows that the behavior of the DMN may be more nuanced than what was earlier believed. We wanted to investigate the dynamics of DMN further and utilized novel statistical approaches to study the same. We analysed four publicly available datasets (total 223 subjects) across seven tasks related to different cognitive modalities and found out that there is a large amount of variability in the presumed task-negative network across subjects, trials and regions. We designed a method called Inter Trial Temporal Synchronization Analysis (IT-TSA) and Inter Subject TSA (IS-TSA) to analyse variability across trials and subjects respectively. We found out that all task-negative regions and even weak task-positive regions have low synchronization across trials and subjects. We hypothesize that the DMN variability might be more due to neural differences rather than structural or HRF variations. We also investigated the use of non GLM technique like the Inter Subject Correlation (ISC) and found that it may only be suitable to study tightly synchronized task-positive regions. Our study challenges the understanding of DMN as a task-negative region and advances the findings that DMN acts as an active component associated with a unique set of self-referential, autobiographical processes which are deactivated differentially and non linearly across trials and subjects in the presence of extrinsic processes.

**Highlights:**

- Default Mode Network is weakly synchronized across trials and subjects in task paradigms.
- Inter Subject Correlation is not suited to detect task-based activations in DMN.
- Synchronization Analysis reveals temporal structure in fMRI data.

## Introduction

The discovery of default mode network was one of the significant findings by cognitive neuroscientists in the past few decades. The accidental observation [1; 2] made while examining the Positron Emission Tomography (PET) images of subjects engaged in externally focused, attention demanding tasks, found that the tasks reduced the activities of several brain regions when compared with a passive condition [3]. Since then a series of studies using task-based fMRI [4; 5; 6] PET [7; 8], resting state fMRI [9; 7; 4] and more recently using the direct neuronal recordings of the local field potentials (LFPs) [10; 11] have confirmed these findings using a variety of task conditions and experimental paradigms. Quite surprisingly, these studies found that the same brain regions consistently show a task-induced reduction of activity irrespective of the task, as long as the task is externally-focused and attention demanding (see [12; 13] for reviews). These regions comprised of the frontal midline, posterior cingulate cortex, inferior parietal lobule, and medial temporal lobe [14] together are now called the default mode network (DMN) of the brain, indicating that these regions are activated in the “default mode” of brain function when there is no externally-focused, attention-demanding task at hand.

Early studies on functional connectivity analysis of resting state fMRI have revealed two anticorrelated brain networks [15; 16] working in tandem with each other. One of the networks, called the task-negative network is the default mode network [17], while the other network, which is negatively correlated to the DMN in resting state comprises of the superior parietal lobule, intraparietal sulcus, frontal eye fields, and ventral premotor cortex and is called the Dorsal Attention Network (DAN). The networks themselves are strongly correlated within and anticorrelated to each other [15]. The DAN is involved in attention demanding extrinsic tasks whereas the DMN is found to be activated during internal self-referential processing [18]. Such anticorrelations were found to be affected by cognitive states, changed across default subsystems and affected by preprocessing methods [19]

Although believed to be task-negative, studies have shown that the behavior of the DMN is more nuanced [6; 17]. Deactivations in the DMN increased with task difficulty [20] and task demands [21]. Task complexity results in gamma suppressions in the DMN regions when investigated using intracranial EEG in a cohort of epileptic patients [22]. DMN activity is predictive of errors [23]. A study analysing the default mode of cats found that the anticorrelations between the DMN and the DAN were found only 20% of the time suggesting a role of DMN in attention [24]. People with disorders like ADHD, Parkinson*’*s disease, Alzheimer*’*s have shown to have differential connectivity in the DMN which can be altered using drugs [25]. The recent use of continuous attention tasks demonstrated a complex dynamics between DAN-DMN activity with task load and attentional control [26]. Another study found variable dynamics of the DMN-DAN interactions which are altered by the frontoparietal control network (FPCN) [19]. Studies that use naturalistic stimuli to study patterns of activity in the brain [27; 28; 29] using methods like Intersubject correlation(ISC) and Intersubject functional correlations(ISFC) have found that the default mode network reconfigures from task-positive to task-negative during naturalistic stimuli. Internal mentation and external monitoring are two leading hypotheses that can be used to describe such a nuanced behavior [30].

Recent work has suggested that DMN may not be a single network but composed of three different subsystems [14; 30]. The exact functions of each of these DMN subsystems is still a subject of active research. Ventromedial PFC reflects the emotional state of the subject whereas the dorsal medial PFC is engaged in self-referential judgements and the activity between both the regions are anti-correlated suggesting difference in the times when these regions are activated [12] The posterior regions of the DMN are more associated with autobiographical memory, emotional and self-referential processing [12; 14]. The MTL subsystem of the DMN is engaged during mnemonic processes, autobiographical memories and recollection based tasks [18]. The posterior cingulate cortex and the anterior mPFC has shown hub like properties as analysed using functional connectivity. Recent studies have shown the involvement of the DMN in task-unconstrained thoughts, mind wandering as well as rumination [14; 31; 32]. Some studies have shown that not only there are multiple networks but these are tightly interwoven across the three large subsystems [33; 34; 13].

In this paper, we test the behavior of Default Mode Network across a variety of tasks and analyse synchronization across trials and subjects. We designed a new method called the Temporal Synchronization Analysis (TSA) which can be applied to multiple trials within a subject or across subjects to determine the synchronization of the BOLD response of different voxels (across trials or subjects) in the brain. We add to the existing body of literature on default mode network and discover that the DMN regions have weaker inter-trial and inter-subject stimulus-locked synchronization as compared to task-positive regions of early sensory processing. These results been have found to be consistent across four very different data sets involving visual-auditory valence, language, craving, risk, emotion, theory of mind and texture matching tasks. Our results demonstrate that the complexity of the dynamics of regions like DMN is not strictly task-negative as have been believed since the last two decades and needs to be studied using newer statistical methods. Inter Subject Correlation (ISC) methods may not be best suited for task-based stimulus-locked paradigms to study the task-negative networks. Given the role of DMN in mind-wandering [14], our method can be used to create a metric to quantify the stimulus-locked DMN synchronization which may be related to the quality of attention control in individuals and predict performance in cognitive tasks across different populations.

## Results

We have detected a weak inter-subject synchronization in the brain*’*s default mode network (DMN) across seven tasks from four different datasets. The Inter Subject Correlation (ISC) [27; 28; 35] method, which finds voxels with large (inter-subject) correlated BOLD activity when applied to a dataset where the task conditions for all the subjects are perfectly synchronized, is expected to find task-positive as well as task-negative regions (in the task-negative regions, the BOLD signal is expected to decrease for all the subjects after the stimulus onset, leading to large inter-subject correlations, see Box 1 for details of the ISC technique). A clear trend can be seen in the scatter plot of GLM zstat values and ISC correlation values of the voxels in the three datasets studied. In Fig. 1(a), the task-negative voxels with very significant (negative) zstat scores generally tend to have low inter-subject correlation values unlike most of the task-positive voxels which exhibit higher inter-subject correlations with higher zstat values. When we applied ISC to the FNF dataset, we found high inter-subject correlations (M=0.19, SD=0.16) in the BOLD signal of early sensory processing regions which were significantly higher [independent samples t-test t(36215) = 127.25, p<0.00001, Cohen*’*s d=1.34] than the task-negative default mode network regions (M=0.03, SD=0.018). ISC for AV dataset task-positive regions (M=0.11, SD=0.07) was significantly higher [t(19877)=129.62, p<0.00001, Cohen*’*s d=1.86] than default mode network regions (M=0.015, SD=0.12) and for the BW dataset, the task-positive inter subject correlations (M=0.072, SD=0.05) were higher but with a medium effect size [t(31206)=6.336, p<0.00001, Cohen*’*s d=0.25] than the default mode network network ISC (M=0.05, SD=0.02). The time series plots in Fig. 1(b)-(e) of a single subject (and subject averages) of task-positive voxel and task-negative voxel shows that there is a reasonable signal change in task-positive as well as task-negative regions during the FNF task. Analysing the four tasks from the HCP dataset, we find a similar pattern of inter-subject correlations as shown in Fig. 2, low for the task-negative regions and significantly higher [Social: t(39623) = 120.13, p<0.00001, Cohen*’*s d=0.79; Emotion: t(24982)=95.23, p<0.00001, Cohen*’*s d=0.87; Gambling: t(38768)=88.41, p<0.00001, Cohen*’*s d=0.84; Relational: t(37839)=106.01, p<0.00001, Cohen*’*s d=0.85] for the task-positive regions.

**Figure 1.**
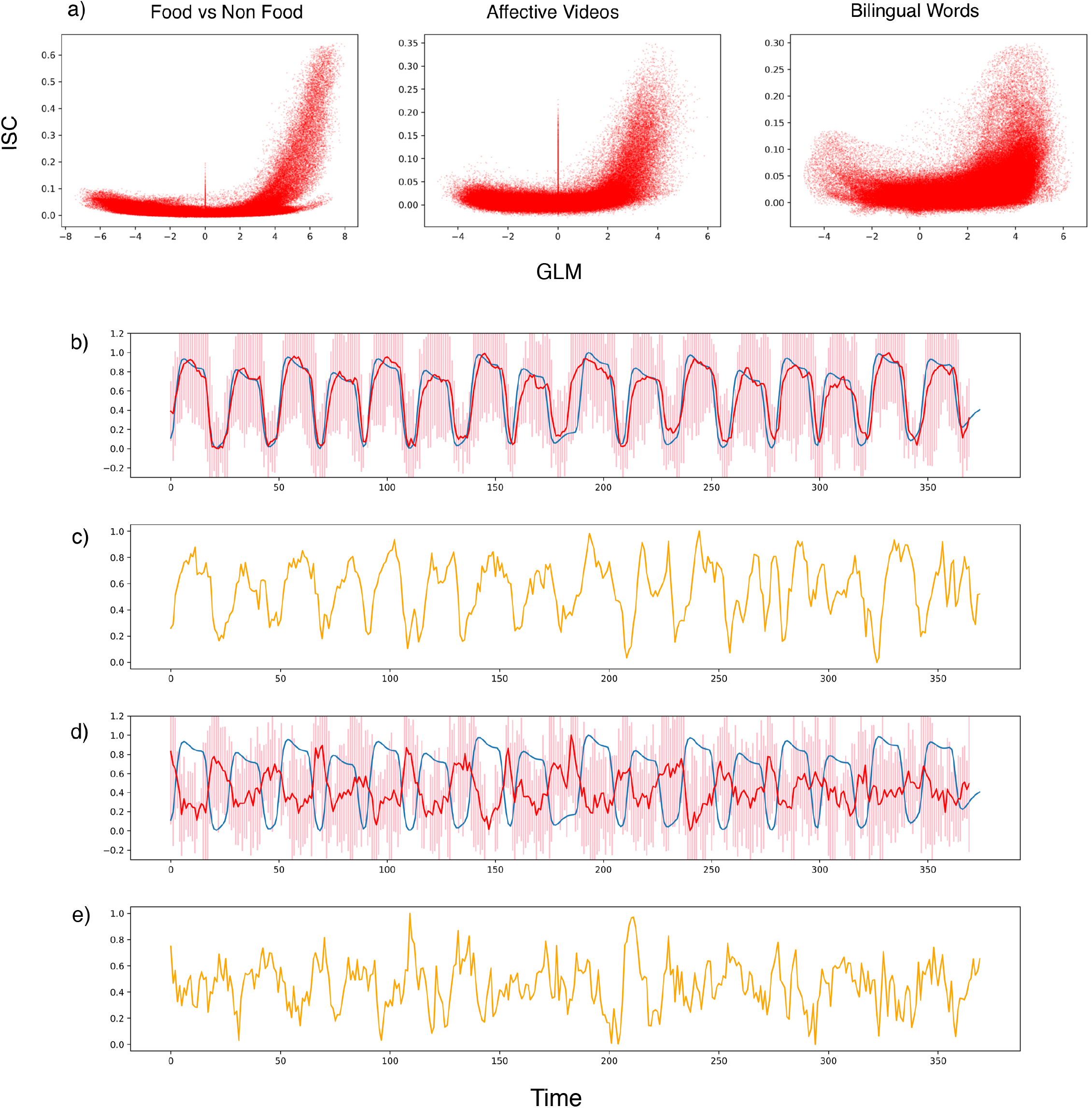
a) A scatter plot of Intersubject correlation values vs GLM zstat values for the three datasets: FNF, AV, BW. b) Blue line represents the design matrix of block-based design of dataset FNF; Red line represents the averaged percent BOLD signal change for voxel (55,16,34) with maximum GLM zstat value and standard deviation as error bar across subjects. c) Percent BOLD signal change for a single subject for voxel (55,16,34). d) Blue line represents the design matrix of block-based design of dataset FNF, Red line represents the averaged percent BOLD signal change for voxel (41,27,47) with minimum GLM zstat value and standard deviation as error bar across subjects. e) Percent BOLD signal change for a single subject for voxel (41,27,47).

**Figure 2.**
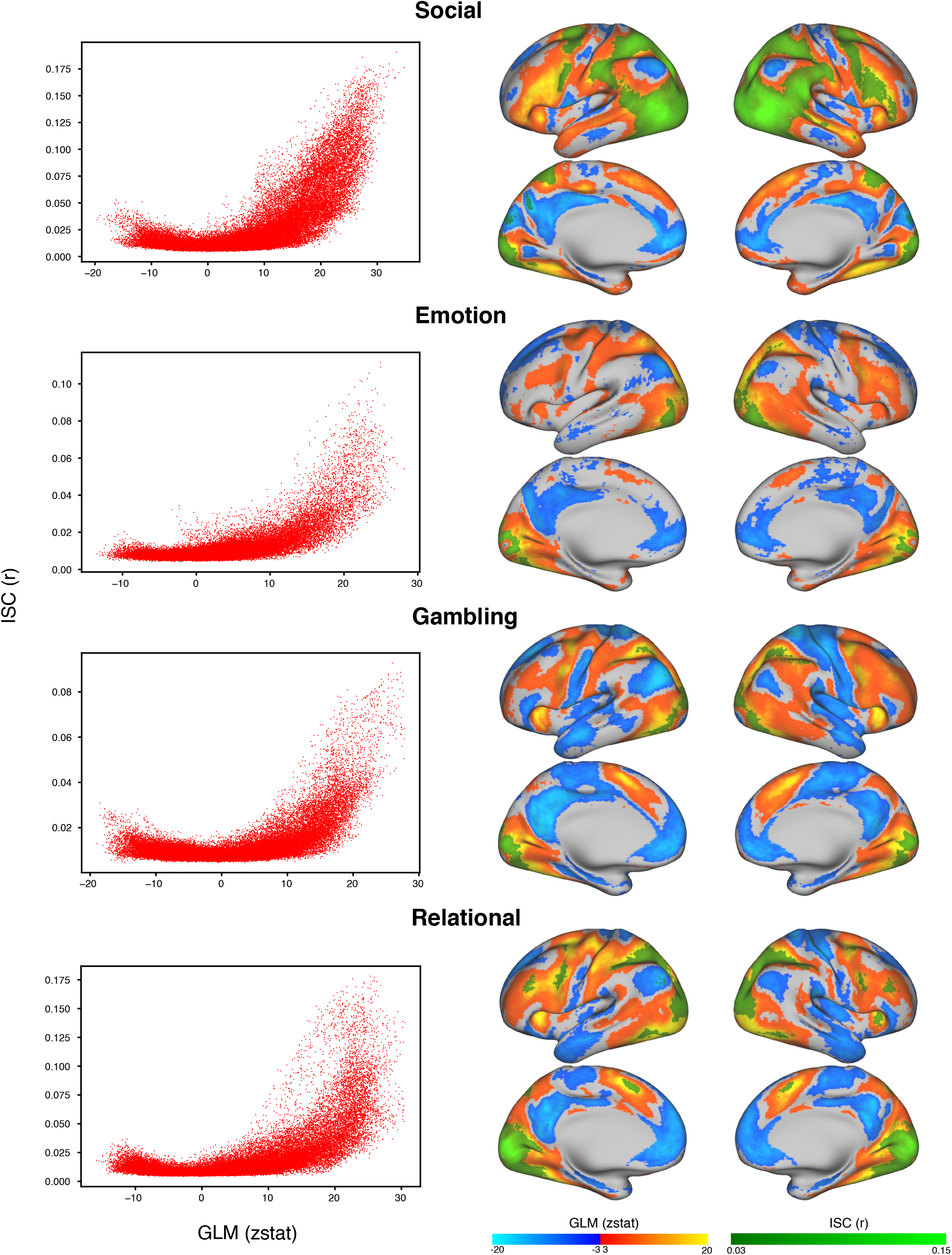
Left panels depict the scatter plots of Intersubject correlation values per vertex on the surface map against the GLM zstat values for the four HCP tasks: Social, Emotion, Gambling and Relational. The right panels are the surface maps for the same.

We can compare the GLM activation maps from the three datasets (AV, FNF, BW) studied with the ISC correlation maps in Fig. 3. There is a good overlap between GLM task-positive voxels and the voxels with significantly high inter-subject correlations in the early sensory processing regions. This overlap is much reduced in the higher-order processing regions. There is nearly zero overlap between the GLM task-negative regions of the DMN and ISC correlation maps. Table 1 quantifies the amount of overlap (using metrics defined in the methods section) between active voxels of GLM and significantly correlated voxels of ISC for different brain regions. Apart from the BW dataset, there is no overlap between the task-negative regions and the ISC correlated voxels. For the task-positive regions there is 33-37% overlap in FNF, 22-23 % overlap in AV and around 61% overlap in BW.

**Table 1.** Overlap between ISC and GLM. M1, M2, M3 and M4 are measures defined in the Methods section. TP - Number of task-positive voxels, TN - Number of task-negative voxels, ISC - Number of activated ISC voxels

| Brain Area | *Dataset* | M1 | M2 | M3 | DICE | TP | TN | ISC |
| --- | --- | --- | --- | --- | --- | --- | --- | --- |
| **Whole Brain** | *FNF* | 0 | 0.078 | 1 | 0.083 | 69104 | 59885 | 5416 |
|  | *AV* | 0 | 0.071 | 0.98 | 0.058 | 37335 | 53988 | 2714 |
|  | *BW* | 0.333 | 0.435 | 0.912 | 0.821 | 132013 | 11174 | 67136 |
| **V1** | *FNF* | 0 | 0.331 | 1 | 0.344 | 5322 | 4910 | 1764 |
|  | *AV* | 0 | 0.235 | 0.979 | 0.333 | 3270 | 1328 | 786 |
|  | *BW* | 0 | 0.629 | 0.903 | 1.169 | 10274 | 100 | 7167 |
| **V2** | *FNF* | 0 | 0.372 | 1 | 0.375 | 6626 | 6521 | 2468 |
|  | *AV* | 0 | 0.223 | 0.981 | 0.325 | 4705 | 1742 | 1072 |
|  | *BW* | 0 | 0.617 | 0.918 | 1.145 | 12358 | 297 | 8304 |
| **Auditory** | *FNF* | 0 | 0 | 0 | 0 | 56 | 1166 | 0 |
|  | *AV* | 0 | 0.197 | 1 | 0.391 | 1364 | 11 | 269 |
|  | *BW* | 1 | 0.424 | 0.543 | 0.627 | 1206 | 2 | 945 |
| **Precuneus** | *FNF* | 0 | 0 | 0 | 0 | 2094 | 11491 | 0 |
|  | *AV* | 0 | 0 | 0 | 0 | 1725 | 4887 | 0 |
|  | *BW* | 0.358 | 0.323 | 0.710 | 0.590 | 5741 | 3006 | 4135 |
| **PCC** | *FNF* | 0 | 0 | 0 | 0 | 30 | 3677 | 0 |
|  | *AV* | 0 | 0 | 0 | 0 | 200 | 2166 | 0 |
|  | *BW* | 0.528 | 0.299 | 0.674 | 0.745 | 852 | 1799 | 1786 |
| **Hippocampus** | *FNF* | 0 | 0 | 0 | 0 | 2258 | 191 | 0 |
|  | *AV* | 0 | 0 | 0 | 0 | 423 | 254 | 0 |
|  | *BW* | 0 | 0.071 | 0.967 | 0.142 | 2463 | 0 | 182 |
| **Paracingulate** | *FNF* | 0 | 0 | 0 | 0 | 903 | 5309 | 0 |
|  | *AV* | 0 | 0 | 0 | 0 | 158 | 5075 | 0 |
|  | *BW* | 0.680 | 0.474 | 0.959 | 1.077 | 4427 | 2649 | 4068 |

**Figure 3.**
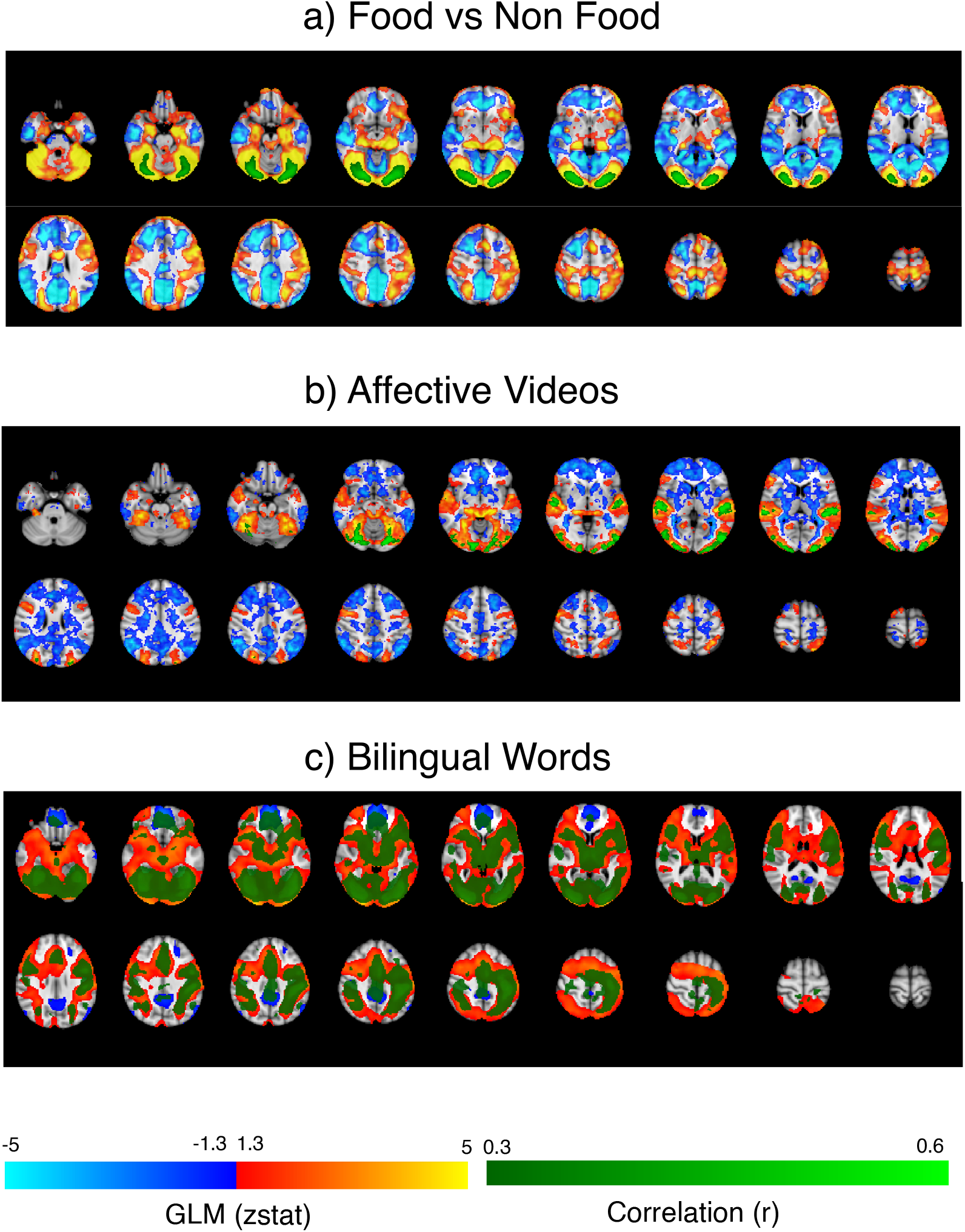
Brain maps of the three datasets showing GLM based activations in red/blue and ISC activations in green.

Why do task-negative regions (and some later task-positive regions), despite having very significant z stats values, do not show any significant inter-subject correlations? It is generally believed that the task-negative regions (primarily comprising the default mode network regions) get deactivated during the presentation of attention demanding stimulus [4]. However, the complete picture is not as straightforward. It turns out that the task-negative regions get deactivated “on the average” for the duration of the task, but such deactivation may not be strictly stimulus-locked. There may be a pattern in such deactivation which may differ from subject to subject and trial to trial. Different subjects (and trials) may get deactivated at different times, decrease in signal may be followed by a subsequent increase in the signal for some subjects (and/or trials) while maintaining an average negative signal change for the duration of the task block. Thus, the GLM analysis which considers the data for all the blocks and all the time-points within these blocks together to estimate the parameter value (beta value) which is then tested for significance, does not consider the dynamics of the BOLD signal within the block. Thus, if a voxel*’*s BOLD response shows a 1% increase followed by a 2% decrease of the same duration, it is likely to be classified as a task-negative voxel that gets deactivated in response to the experimental condition by the conventional GLM analysis.

### Box 1.

**Inter Subject Correlation**

**Box 1 Figure 1.**
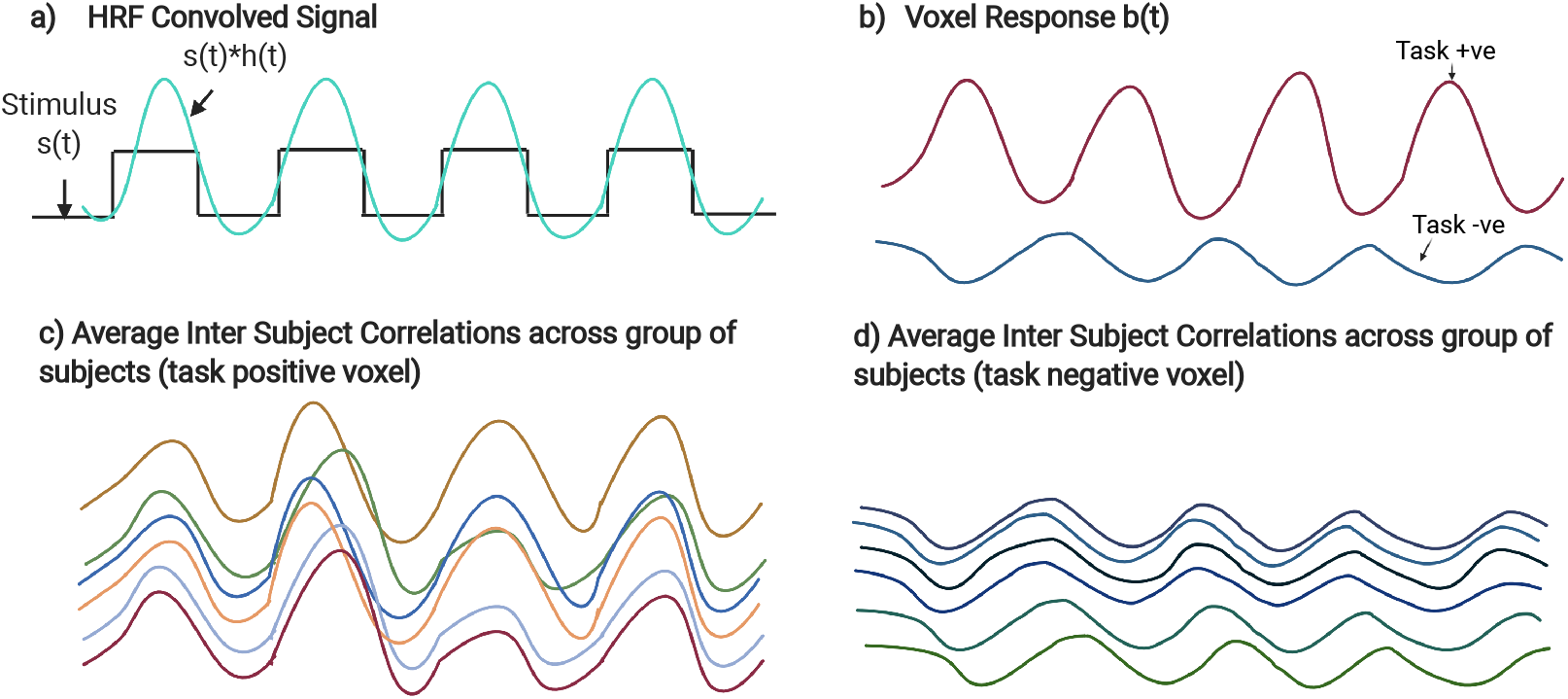
ISC: a) ISC method applied to block-based task paradigms. b) Voxel responses for task-positive and negative region differ and result in different ISC values across subjects in c) and d).

The Inter-subject Correlation (ISC) method is a promising approach for statistical analysis of fMRI data [27]. Although it has been claimed that ISC may be used to *fi*nd brain activations in the conventional block-based or event-based (or mixed) task paradigms [35], with good concordance to the General Linear Model (GLM) analysis results, the unique value of the ISC method comes from its unique ability to *fi*nd brain activations in naturalistic paradigms such as watching a movie inside the fMRI scanner [36], listening to audio stories [37] or other similar paradigms [29]. The conventional GLM analysis method, which requires an explicit definition of different task conditions, is too restrictive in such paradigms as compared to the ISC method where no such definitions are needed.

In the experimental paradigms suitable for the ISC analysis, all the subjects are given the same stimulus in the fMRI scanner. The BOLD time series of a voxel is then correlated with the BOLD time series of the corresponding voxels of all the other subjects. Statistical techniques such as bootstrapping [35] are used to find voxels with significant positive correlations with the corresponding voxels of other subjects.

A significant Inter Subject Correlation in a voxel*’*s BOLD time series indicates that the corresponding voxels of the subjects are responding similarily to different experimental conditions i.e., they must be getting activated and de-activated together. Since the subjects are scanned independently and they only share identical experimental conditions, it may be concluded that the simultaneous activations and de-activations of the voxels of different subjects must be in response to the different experimental conditions presented during the experiment. The ISC method has been very successful in finding brain activity under different types of naturalistic experimental paradigms as it does not require an explicit definition of different experimental conditions.

When the ISC method is applied to a task-based paradigm with identical stimulus timings across the subjects, it is also expected to find significant inter-subject correlations among the task-negative voxels of the default mode network (DMN). At the presentation of an externally-oriented, attention-demanding stimulus, the BOLD response of a DMN voxel of all the subjects is expected to reduce (and is expected to increase when the stimulus is removed) thereby causing a significant inter-subject correlation among them (see Fig. 1(d)). We however find that this is not the case. The lack of synchronization in the DMN deactivations leads to low ISC of voxels in the DMN.

It turns out that the deactivations in the default mode network regions are not strongly locked to the stimulus and are also not strongly synchronized across the trials or the subjects. Fig. 4 shows the average percent signal change with respect to the trial onset for the FNF dataset. Here, we reorganized the data into corresponding trial blocks and plotted the percent BOLD signal change from the block onset and averaged it across trials. The top row shows the data for a single subject and the bottom row shows the data for the average of subjects. As we can see from panels in the first two columns that the trial-average and subject-trial-average deactivations in the default mode network regions have a very low percentage change (M=0.15% as compared to 1%). This is since deactivations in different trials and different subjects occur at different times and averaging this across trials or subjects cancel the effect out leading to an overall lower magnitude of percentage signal change (and also lower inter-subject correlations). The amount of synchronization in task-positive regions is considerably higher as is evident from the task-positive panels in Fig. 4.

**Figure 4.**
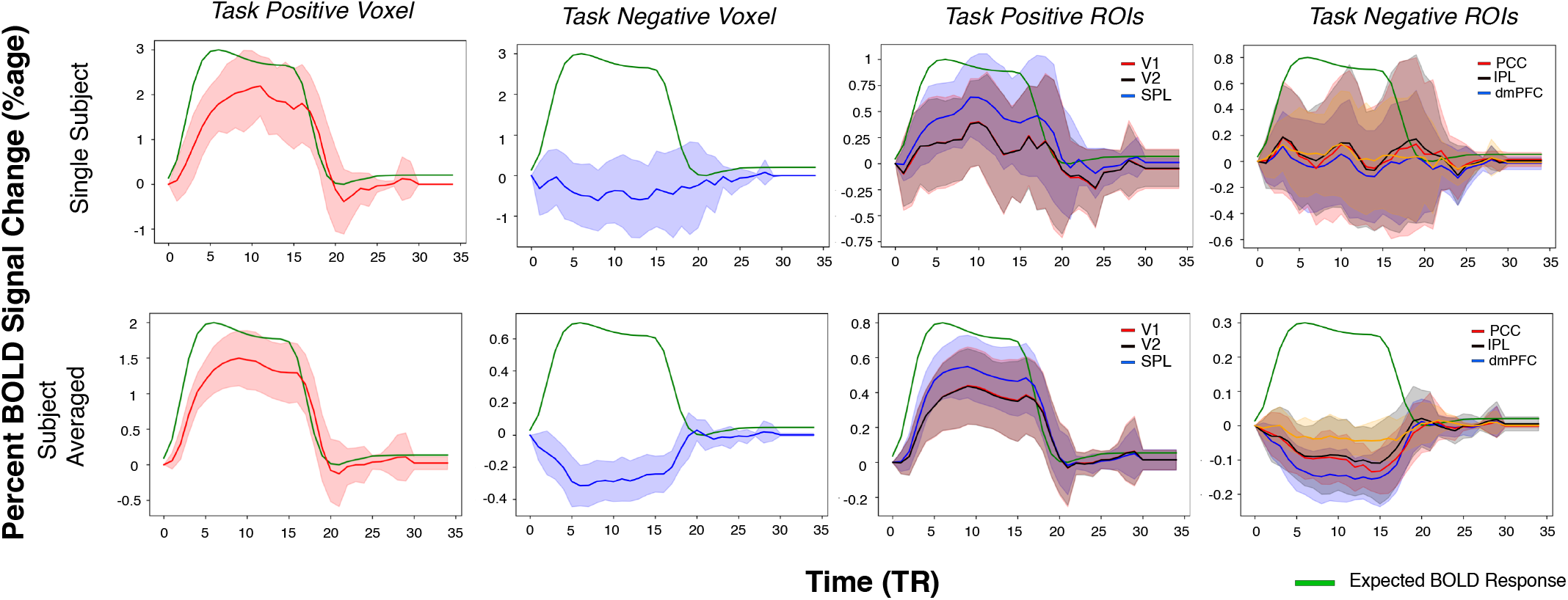
Percent BOLD signal change with respect to time of trial onset averaged across all the blocks in the task. The top row refers to data from a single subject, averaged across the blocks, and the bottom row refers to subjects*’* averaged data. First columns represents data for max zstat voxel (55,16,34), and second columns for min zstat voxel (41,27,47),. The averaged signal change for task-positive: V1, V2, Superior Parietal Lobule and task-negative: Inferior Parietal Lobule, dorsal medial prefrontal cortex and Posterior Cingulate Cortex ROIs are shown in third and fourth columns respectively. The task-positive and negative ROIs were derived from the Schaefer atlas [38]. The green curve depicts the expected BOLD responses within a block. The shaded region depicts the standard error of the mean (SEM) across trials.

To characterize the synchronization, we developed two approaches called Inter Trial Temporal Synchronization Analysis (IT-TSA) and Inter Subject Temporal Synchronization Analysis (IS-TSA) which respectively measures the significance in synchronized BOLD signal change across trials or subjects respectively at each time instant (see Methods for details). Fig. 5 is laid out similar to Fig. 4 and depicts how the IT-TSA zstat value varies with time within a trial interval for an individual task-positive and task-negative voxel (left two columns) and for default mode network and task-positive ROIs (right two columns). For the most task-positive voxel, the absolute IT-TSA zstat at mid of block (M=8.78, SD=3.67) significantly differs [t(29)=10.06, p < 0.0001, Cohen*’*s d = 2.45) with the most task-negative voxel (M=-2.03, SD=1.15) implying much weaker inter-trial synchronization in the default mode network regions. The subject averaged absolute IT-TSA value at mid-block for task-positive ROIs V1 [M=4.65, SD=2.44], V2 [M=5.21, SD=1.76], and Superior Parietal Lobule [0.90, SD=0.7] was higher than default mode network ROIs Posterior Cingulate Cortex (M=1.31, SD=0.26), Inferior Parietal Lobule (M=1.36, SD=0.22), and dorsal medial prefrontal cortex (M=0.98, SD=0.14). We performed an independent t-test to determine the difference between the IT-TSA zstat values for the default mode network and task-positive regions. Fig. 6(a) shows the ROI averaged IT-TSA values for three task-positive (V1, V2, SPL) (combined mean=3.34, SD=2.61) and three default mode network regions (PCC, IPL, dlPFC) (combined mean=1.19, SD=0.27). The task-positive regions were more synchronized across trials [t(10641)=64.86, p <0.0001, Cohen*’*s d=1.15]). Fig. 6(b) shows the plot of ROI averaged IT-TSA values versus GLM Z-statistic. The task-positive regions (averaged GLM zstat > 2) shows higher synchronization than default mode network regions similar to other non task-positive regions (zstat < 2). We fit a simple linear regression to predict IT TSA values based on the GLM zstat values separately for task-positive and non task-positive including the default mode network. For the task-positive regions, significant regression was found [F(1,22)=97.09, p<0.0001, *R*^2^=0.815; IT-TSA=-2.01+1.04*GLM]. IT TSA value increased by 1.04 for every unit increase in the GLM zstat for task-positive regions. For the non task-positive regions also the regression fit was significant [F(1,74)=452.6, p<0.0001. *R*^2^=0.859; IT-TSA=0.021+0.167*GLM]. IT TSA value decreased by 0.167 units for every decrease in one unit of GLM zstat value for non-negative and default mode regions. Fitting the linear regression on only the task-negative regions, we found significant fit [F(1,29)=63.62,p<0.0001, *R*^2^=0.687; IT-TSA=0.19+0.224*GLM]; every unit decrease in GLM zstat decreased the IT TSA by 0.224 units.

**Figure 5.**
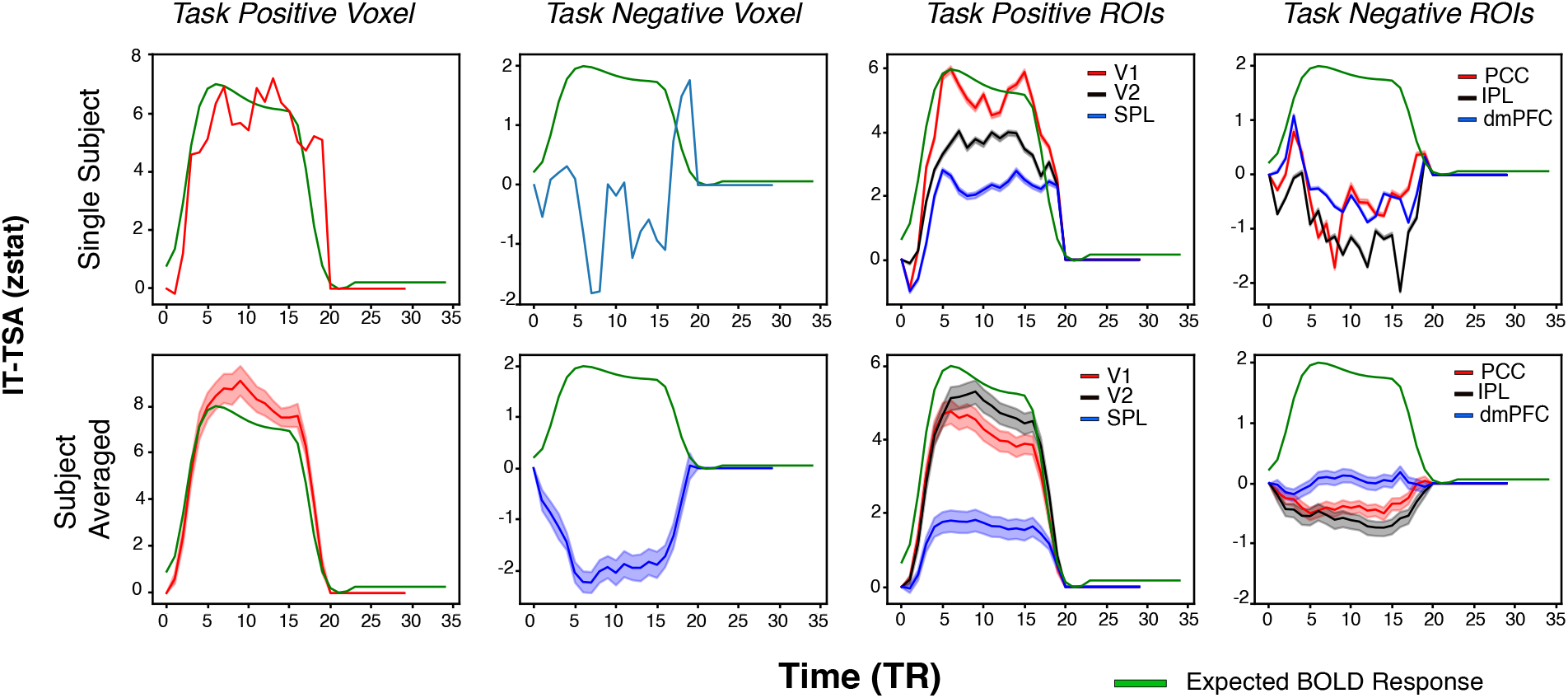
We plotted the Inter Trial Temporal Synchronization Analysis parameter for a single subject and averaged subjects for the FNF dataset. The top row refers to a single subject and the bottom row to averaged subjects. First column is for data from voxel (55,16,34) with max zstat (red), second column is for data from voxel (41,27,47) with minimum zstat value (blue). Third column plots the task-positive ROIs: V1, V2 and Superior Parietal Lobule. The fourth columns plots the task-negative ROIs: Posterior Cingulate Cortex, Inferior Parietal Lobule and dorsal medial Prefrontal Cortex. The green curve depicts the ideal BOLD response. The shaded area in the top row represents the SEM across voxels in ROIs and the bottom row represents SEM across subjects.

**Figure 6.**
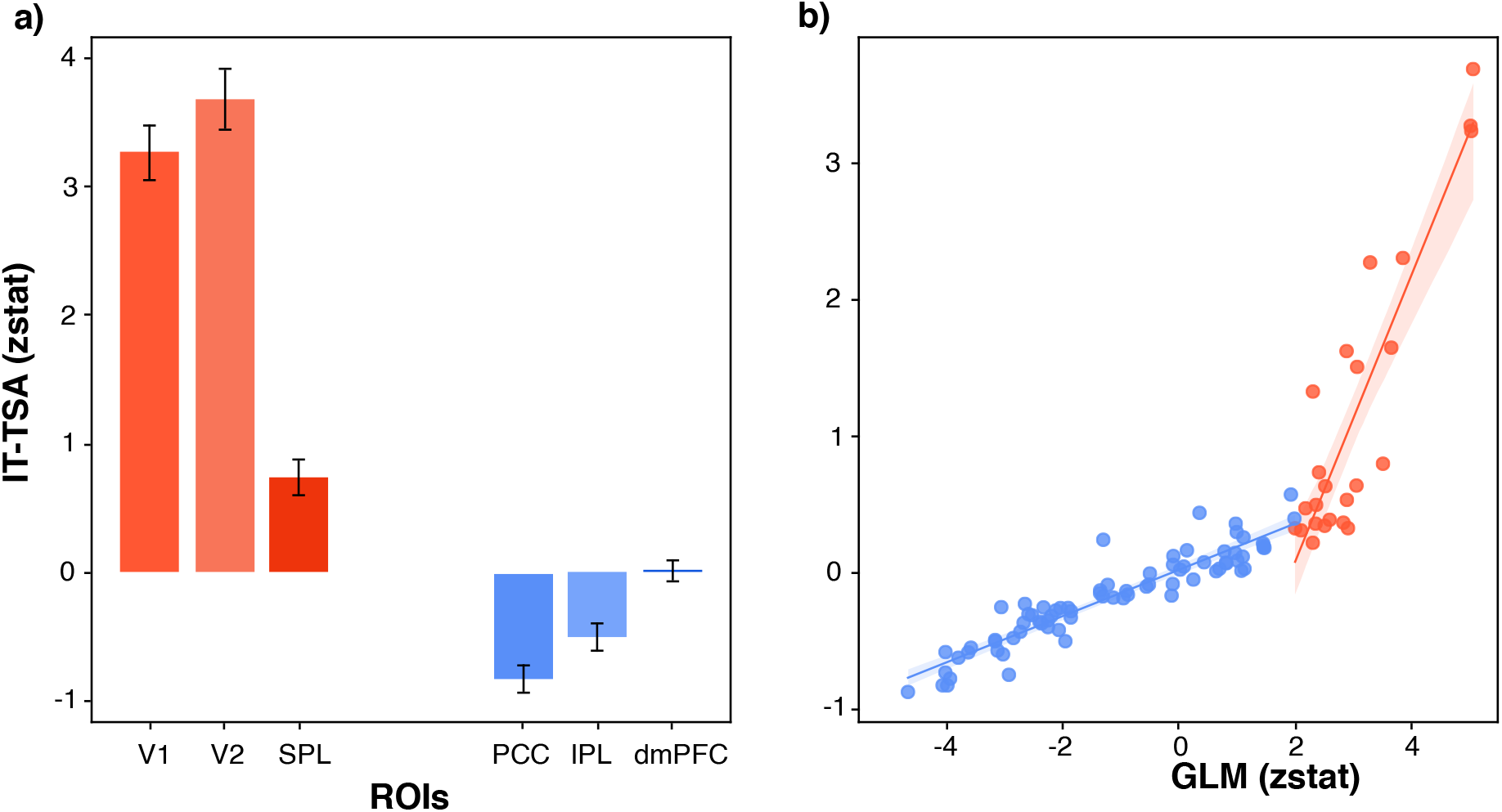
Analysis of the IT-TSA values across ROIs from the Schaefer atlas in the FNF task. The left panel represents the bar plot for task-positive and non task-positive ROIs. The max IT-TSA from each averaged signal across subjects for each voxel was computed. The bar plot represents the meaned IT-TSA across voxels in the corresponding ROI and the standard deviation. The error bar represents SEM across the ROIs. The right panel indicates the scatter plot between the ROI averaged zstat value v/s ROI averaged IT-TSA value. The shaded area represents the CI of curve fit.

To determine synchronization across subjects, we plotted the IS-TSA in Fig. 7 for the FNF dataset and in Fig. 8 for the AV dataset. The most significant task-positive voxel in the FNF dataset had higher absolute IS-TSA (M=7.32, SD=4.78) as compared to the most task-negative voxel (M=1.7, SD=1.33).

**Figure 7.**
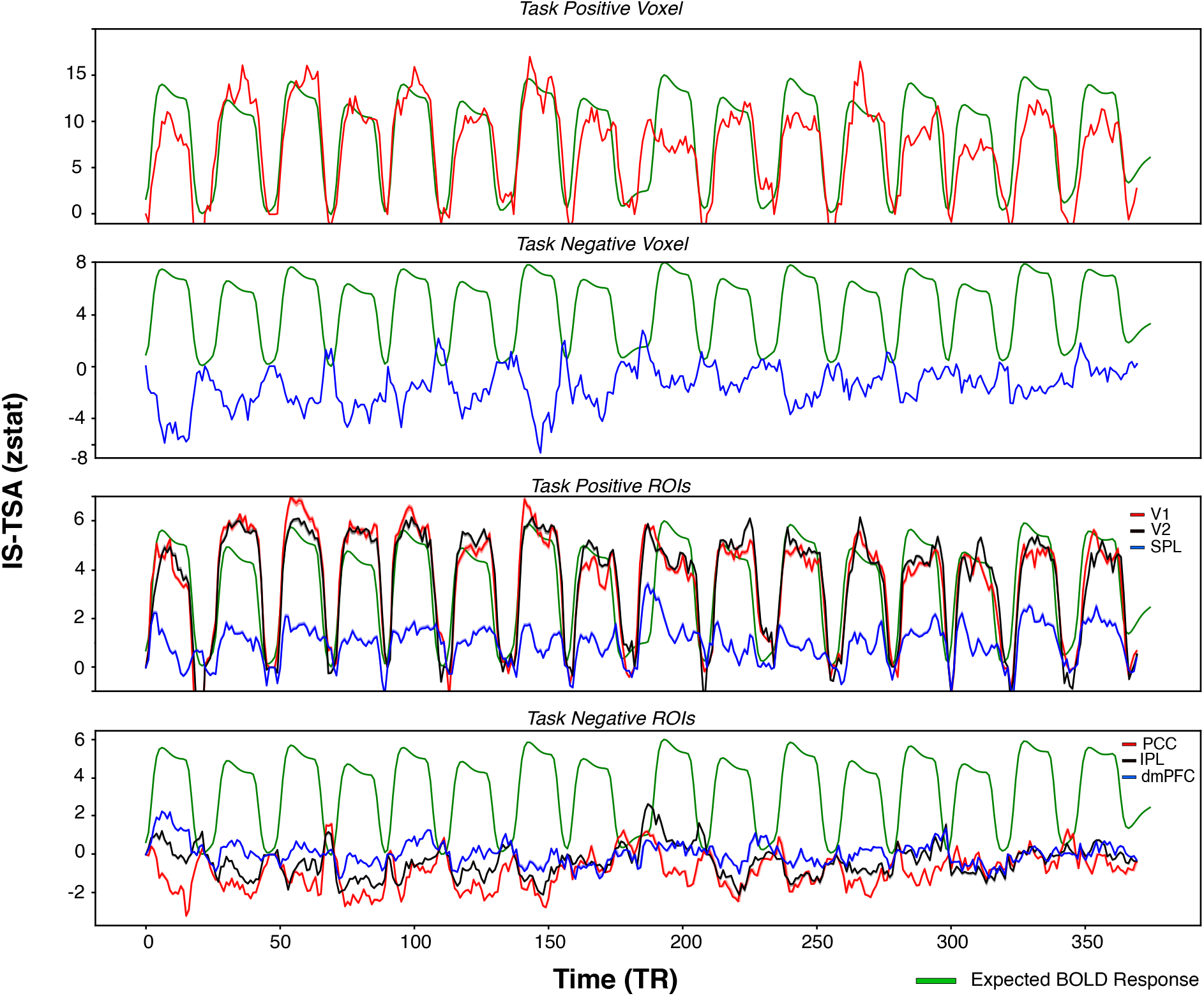
IS-TSA values across time for the FNF dataset. The top panel depicts IS-TSA for a voxel with maximum GLM zstat value, second panel with the minimum GLM zstat value. The bottom two panel shows the averaged IS-TSA across voxels for task-positive ROIs: V1, V2, SPL and default mode network ROIs (PCC, IPL, dmPFC). The green curve depicts the ideal BOLD response. THe shaded region depicts the SEM across ROIs.

**Figure 8.**
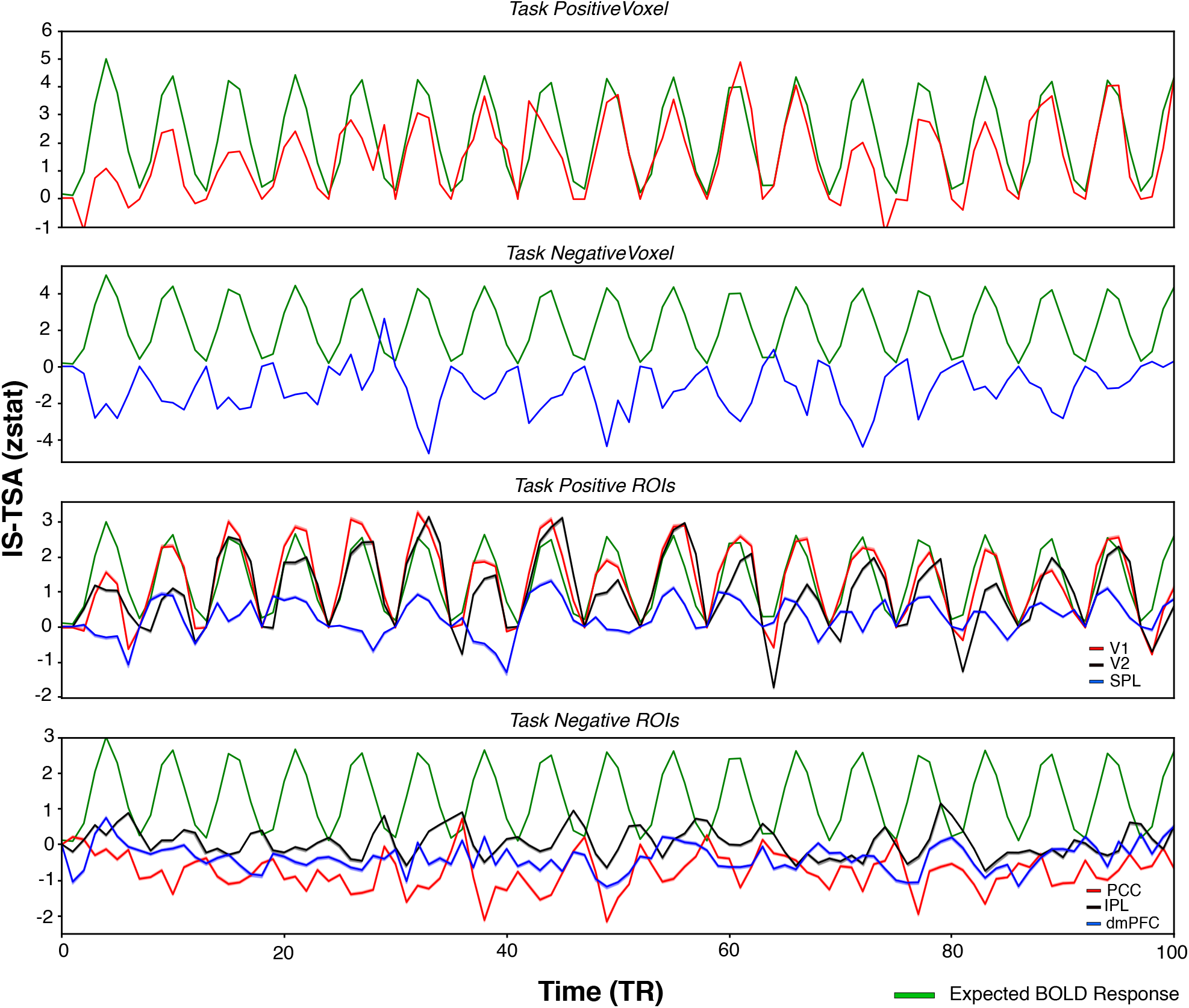
IS-TSA values across time for AV dataset. The top panel depicts IS-TSA for a voxel with maximum Z statistic value, second panel with the minimum Z statistic value. The bottom two panel shows the averaged IS-TSA across voxels for task-positive ROIs (V1, V2, SPL) and default mode network ROIs (PCC, IPL, PFC). The green curve depicts the ideal BOLD response. The shaded region depicts the SEM across ROIs.

For the task-positive ROIs, the IS-TSA values at the middle of a task block for V1 (M=4.93, SD=2.81), V2 (M=5.50, SD=2.34), SPL (M=0.73, SD=1.43) are collectively significantly higher [t(10641)=49.55, p <0.00001, Cohen*’*s d = 0.89) than the absolute IS-TSA values at middle of block for the task-negative ROIs - PCC (M=2.3, SD=1.43), IPL (M=1.44, SD=0.84) and dmPFC(M=0.79, SD=0.54). Similarly for the AV data set the IS-TSA values for task-positive ROIs V1 (M=3.00, SD=1.45), V2 (M=3.47, SD=1.79), SPL (M=1.42, SD=1.06) is significantly higher [t(9000) = 6.25, p <0.0001, Cohen*’*s d=1.205] as compared to absolute IS-TSA values middle of block for default mode regions PCC (M=0.91, SD=0.68), IPL (M=0.99, SD=0.70), and dmPFC (M=0.82, SD=0.61). We averaged the IS-TSA values across the Visual, Dorsal Attention and Default Mode Networks as defined in the Yeo atlas [39] for the four HCP tasks as shown in Fig. 9 and perform independent samples t-test across the networks. We found that visual regions have significantly higher [Social: t(24447)=80.03, p<0.00001, Cohen*’*s d=1.15; Emotion: t(24447)=113.84, p<0.00001, Cohen*’*s d=1.71; Gambling: t(24447)=88.82, p<0.00001, Cohen*’*s d=1.29; Relational: t(24447)=106.24, p<0.00001, Cohen*’*s d=1.53] IS-TSA values across the four tasks as compared to DMN. We saw a similar difference between DAN and DMN [Social: t(24852)=95.35, p<0.00001, Cohen*’*s d=1.31; Emotion: t(24852)=99.88, p<0.00001, Cohen*’*s d=1.38; Gambling: t(24852)=41.50, p<0.00001, Cohen*’*s d=0.55; Relational: t(24852)=70.42, p<0.00001, Cohen*’*s d=0.97]. These results establish that although the default mode network regions do show a BOLD signal decrease during the trial presentation, this signal decrease is weakly synchronized across trials and subjects.

**Figure 9.**
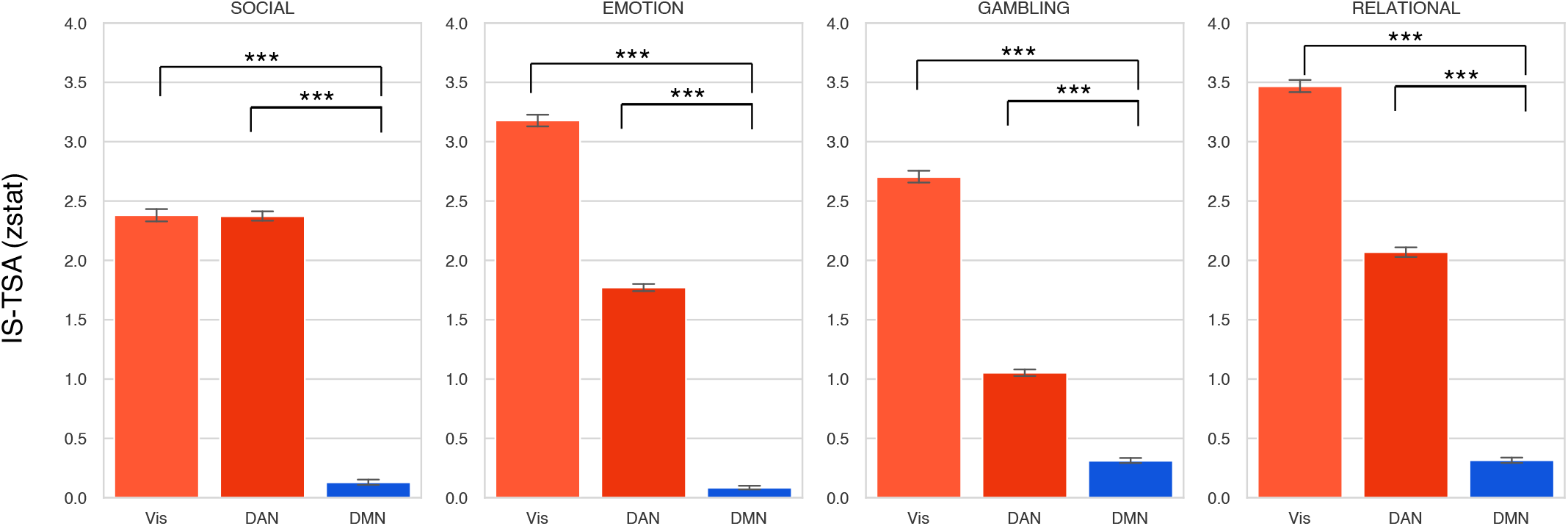
IS-TSA values averaged across the Visual, Dorsal Attention and Default Mode Networks as defined in the Yeo 2011 atlas for the four HCP tasks. Error bars indicate within network standard error of the mean. *** denotes significance p < 0.00001.

We then compared the ROI averaged IS-TSA zstat value against the GLM zstat value for the three tasks in Fig 10(a). The task-positive regions (GLM averaged zstat > 0) show higher IS-TSA values as compared to non task-positive regions and default mode regions. We found a significant regression [F(1,47)=98.33, p<0.0001, *R*^2^=0.677; IS-TSA=-0.75+0.82*GLM] for the task-positive regions, and the IS TSA zstat value increased by 0.82 for every unit increase in the GLM zstat. For the non task-positive regions also the regression fit was significant [F(1,49)=63.56, p<0.0001. *R*^2^=0.56; IS-TSA=-0.11+0.41*GLM]. IS TSA value decreased by 0.41 units for every decrease in one unit of GLM zstat value for non-negative and default mode regions. From the plot in Fig 10(a), we see that the default mode regions have lower IS-TSA values. We fit a linear regression model and found significant regression for task-positive regions [F(1,49) = 94.08, *R*^2^=0.658; IS-TSA=0.30+0.742*GLM] and for task-positive regions [F(1,47]=62.17, p<0.0001, *R*^2^=0.569, IS-TSA=0.43+0.46*GLM] which signifies that for every unit increase in GLM zstat, task-positive regions IS-TSA increased by 0.78 units but only 0.46 units for task-negative regions. For the BW dataset from the plot in Fig 10(a), the linear fit between the IS-TSA and GLM values for task-positive regions [F(1,82)=53.10, p<0.0001, *R*^2^=0.393; IS-TSA=0.22+0.50*GLM] and task-negative regions [F(1,13)=5.196,p<0.05,*R*^2^=0.286;IS-TSA=1.06+0.58*GLM] was significant but there was no difference in the slopes of the regression parameters although the IS-TSA values were higher for task-positive regions [t(7879)=26.51, p<0.00001, Cohen*’*s d = 0.03], the effect size was negligible.

**Figure 10.**
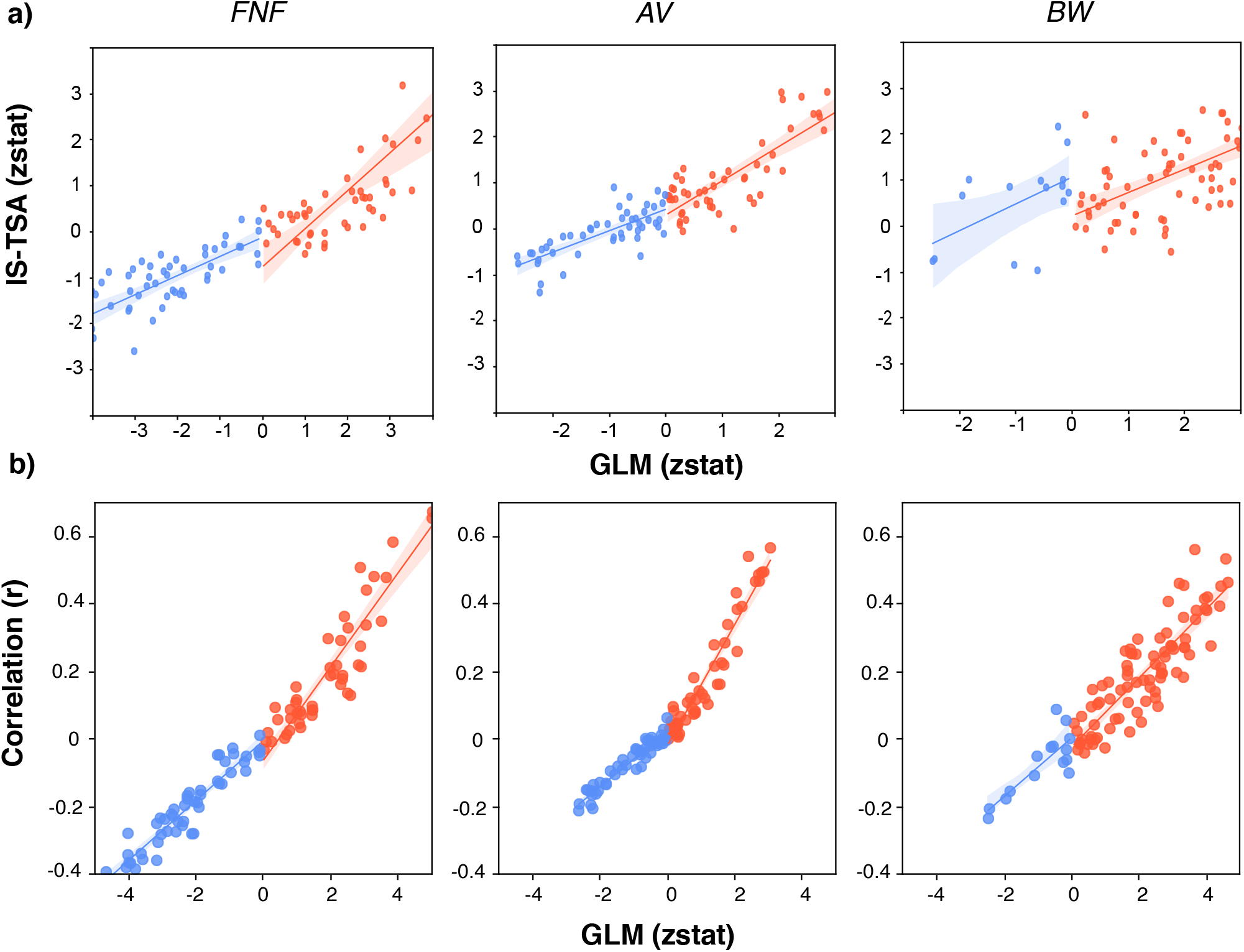
A) IS-TSA value averaged across ROIs from the 100 parcel Schaefer atlas plotted against the ROI averaged GLM zstat values indicating a stronger slope for the task-positive regions than the task-negative regions. B)The IS-TSA value per voxel was computed for the three datasets and correlated with the HRF convolved stimulus signal across time. The correlation was then averaged across all the voxels in the ROI from the Schaefer atlas and plotted with the ROI averaged Z statistic value. IS-TSA values are correlated strongly with the stimulus for task-positive regions than default mode network regions for AV and FNF dataset. The effect is less significant for the BW dataset. The shaded region represents the CI of linear regression fit.

In order to calculate the efficacy of the IS-TSA metric, we plotted the stimulus signal correlated with the IS-TSA for each voxel which was then averaged over the ROIs from Schaefer atlas [38] and plotted for the three datasets in Fig 10(b). The scatter plots were fit to a linear regression model separately for task-positive and default mode network regions for the three datasets. The statistics for the linear regressions [F(1,47) = 286.2, *R*^2^=0.859; IS-TSA=-0.06+0.138*GLM for task-positive, CI=(0.122,0.155); F(1,48) = 373.2, *R*^2^=0.886; IS-TSA=-0.005+0.08*GLM, CI=(0.079,0.098) for task-negative regions] for the FNF dataset show that stimulus correlation with the IS-TSA increases more by 0.06 per unit change in GLM zstat, similarly [F(1,48) = 570.0, *R*^2^=0.922; IS-TSA=-0.02+0.179*GLM, CI=(0.164,0.194) - task-positive; F(1,47) = 529.9, *R*^2^=0.919; IS-TSA=0.02+0.081*GLM, CI=(0.074,0.089) - task-negative] by 0.1 for AV dataset and 0.02 [F(1,82) = 243.7, *R*^2^=0.748; IS-TSA=-0.02+0.102*GLM, CI=(0.089,0.115) - task-positive; F(1,13) = 27.32, *R*^2^=0.678; IS-TSA=0.005+0.086*GLM, CI=(0.051,0.122) - task-negative] for BW dataset, indicating IS-TSA values are less synchronized and relatively weaker for the default mode network as compared to the task-positive regions.

## Discussion

In this paper, we analyzed the synchronization of stimulus-locked deactivations in the default mode network (DMN) and found that the task-induced deactivations in DMN regions have a weaker synchronization across trials and subjects as compared to synchronization in stimulus-locked activations in task-positive regions. This effect is consistent in seven vastly different tasks involving varied regions of the brain. There may be two plausible explanations for this observed phenomenon: *Physiological* or *Neural*.

### Physiological explanation

The fMRI BOLD signal measures the changes in the blood oxygenation level (which is related to the cerebral blood flow, CBF), which is indirectly related to the neural activity through an empirically observed *hemodynamic response function* (HRF) (see Box 2). The exact mechanism of this neurovascular coupling is still not very well understood and is a topic of active research [10; 40; 41]. Many early studies on the HRF have reported (though slightly inaccurately – see (Box 2) a significant variability in the HRF across voxels [42; 43] and regions of interest (ROIs) [44; 45], subjects [46], trials [17] and experiments [47]. The lack of inter-trial, inter-subject and inter-voxel synchronization in DMN deactivations may be attributed to the corresponding variability in the HRFs. The possible reasons for the (incorrectly) observed variability in the HRF as discussed in the literature are differences in vasculature [44], duration of stimulus [47; 48; 49; 41], presentation rate [42], laminar differences [50; 51; 52], ventricle size, density, and vessel elasticity [53]. However, the above reasons do not sufficiently explain why weaker synchronization is observed only in the DMN regions, not in most of the task-positive regions.

### Neural explanation

The observed BOLD signal is the convolution of the HRF and the neural signal (see Box 1). The weaker synchronization of the BOLD signal in the DMN regions may be due to the weaker synchronization in the corresponding neural signal (or neural activity). For this explanation, one needs to first examine the functions of DMN which is still an active area of research [13; 14; 31]. Presently two predominant hypotheses (namely, the *Sentinel Hypothesis* and the *Internal Mentation Hypothesis* [30] have been proposed to explain the observed task-induced deactivations in the DMN regions, both of which can very well explain the observed weaker stimulus-locked synchronization in the DMN regions.

According to the sentinel hypothesis, the DMN plays a role in monitoring the external environment [7; 3] and when presented with an active task requiring focused attention, the brain directs its inner resources to attending the task, while temporarily suspending the environment monitoring. The “default network is hypothesized to support a broad low-level focus of attention when one – like a sentinel – monitors the external world for unexpected events” [3; 54]. If the sentinel hypothesis is true, it may be argued that different trials and different subjects take different amounts of time to disengage from the external monitoring and start attending to the presented task thereby leading to the weaker task-locked synchronization in the neural signal.

According to the internal mentation hypothesis, the DMN directly contributes to internal mentation such as self-reflective thoughts and judgements [18; 55]. Imaginative constructions of hypothetical events or scenarios [56], autobiographical recall [57], theory of mind related activity [58]. When presented with an attention demanding task, the internal mentation is temporarily suspended to attend to the task. Under this hypothesis also, different trials and subjects may take different amounts of time to *disengage* from their internal mentation and attend to the task at hand thereby leading to variability in the neural signal of the DMN. As a result, the observed BOLD signal is expected to have weaker synchronization as compared to the task-positive regions, especially those involving the early sensory processing.

In addition, there is more evidence favouring the neural hypothesis for explaining the weaker synchronization in the DMN regions. Some studies demonstrate the dynamic reconfiguration of the DMN regions indicating that the neural activity in different DMN regions may not always be positively correlated and the activity of some DMN regions may not always be negatively correlated with task-positive regions. Some studies report changes in the function of DMN with different disease conditions.

#### Interdigitated Networks

Recent work by Braga and colleagues [33; 60] have shown that DMN regions are interspersed and juxtaposed networks which are hard to parse out in group averaged results and may slightly vary across subjects which would affect any technique performing group level analyses.

#### Dynamic Reconiguration of DMN

Though Default Mode Network deactivations are related to the task dificulty [6], engagement and the activity in sensory cortices are related to signal suppression in the DMN [17] but recent evidence have found that DMN may dynamically reconfigure with the task. A study looked at the simultaneous EEG-fMRI data in a visual oddball paradigm and found transient engagement in both task-dependent and default mode networks on a millisecond timescale [61]. Such a reconfiguration was also found in the default networks when subjects watched a movie in the fMRI scanner [29]. During a decision making task, the deactivation of the DMN did not occur for all subjects and reduced deactivations were not related to impaired task performance [62]. In a gradual onset continuous performance task(gradCPT), the authors found an associated role of DMN-DAN regions in maintaining attention and trial by trial variability in the activations of task-positive regions and deactivations of the DMN region was related to prestimulus alpha power [63].

#### Effect of Disorders on DMN

Various brain disorders can affect the default mode networks. Mohan et. al.(2016) summarises the effects of Parkinson*’*s, Alzheimer*’*s and Attention Deficit Hyperactivity Disorder (ADHD) on the functional connectivity and how using drugs like memantine and donepezil can restore the functional connectivity in cases of Alzheimer*’*s [25]. Adolescents with ADHD show higher hemodynamic response variability possibly due to neural fluctuations in the anterior regions of DMN as compared to healthy controls [64] which are associated with reduced task performance [65]. Patients with ADHD are unable to suppress their DMN [66] which may have dysfunctional interactions with the executive control network [67]. A study found that the synchronization between task-positive network and DMN fails in case of ADHD and when restored using the drug methylphenidate improved task performance [68].

##### Box 2.

**BOLD Variability**

**Box 2 Figure 1.**
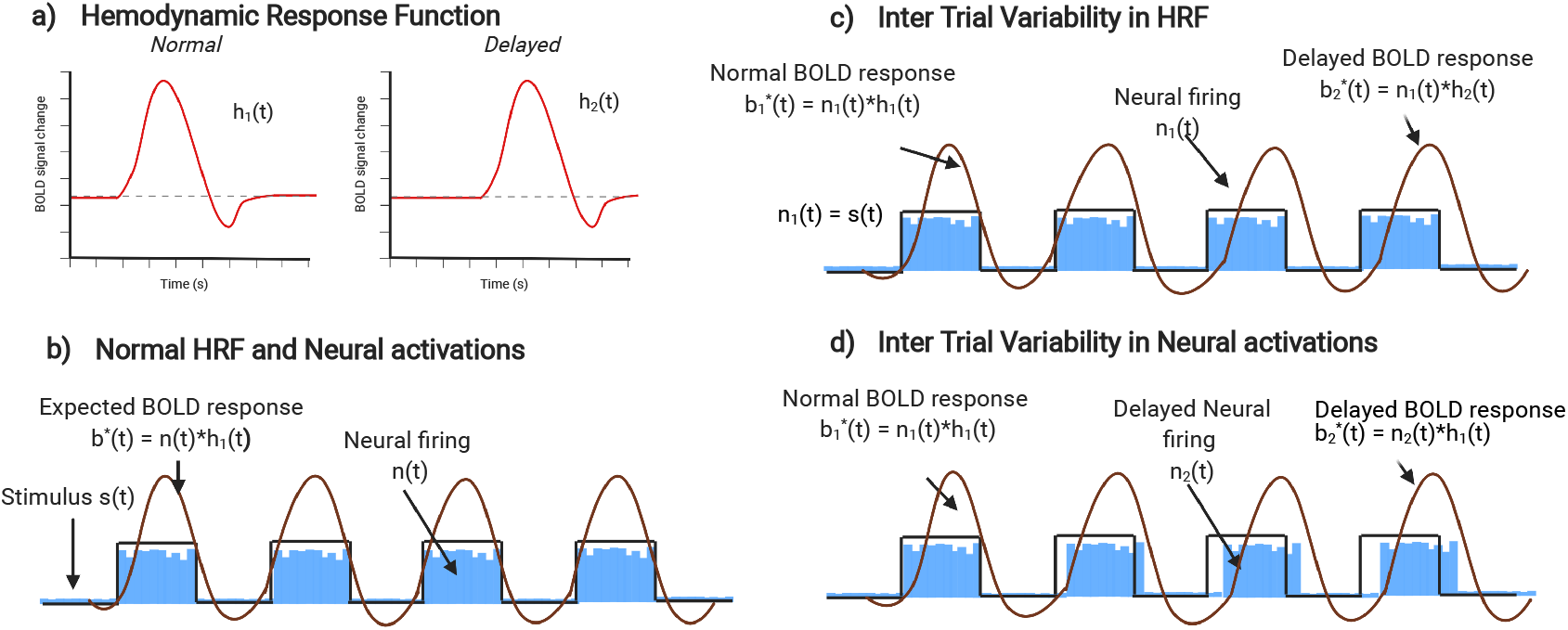
We look at the BOLD signal variability caused by delayed HRF or delayed neural firing. a) Normal and delayed HRF. b) Normal HRF and neural signals result in a stimulus locked expected BOLD response. c) Delay in HRF within trials can result in a delayed BOLD response in later trials. d) The same BOLD response can be detected if the neural firing gets delayed as trials progress.

General Linear Model:

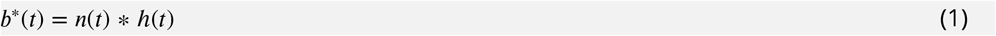

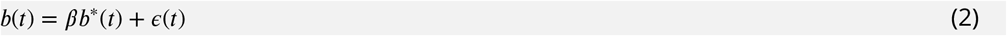

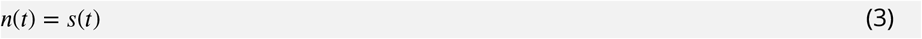

- *b*(*t*): The observed BOLD signal
- *b*^*^(*t*): Expected BOLD response
- *h*(*t*): The hemodynamic response function (HRF)
- *n*(*t*): The neural signal (representing the neural activity)
- *s*(*t*): The activating stimulus
- *e*(*t*): Noise

The observed BOLD signal of a voxel in the area of activity has been empirically observed to be delayed and seems to have gone through a low pass filtering (see Fig. 2(a)). Most of the fMRI analysis models it as a linear-time invariant (LTI) system and the corresponding transfer function mapping the experimental condition to the ideal BOLD response is called the hemodynamic response function (HRF). Another implicit (and potentially incorrect) assumption made in most of the fMRI modeling literature, especially those which study the variability in the HRF (such as [47; 59; 46; 48; 43]) is that the neural response *n*(*t*) is identical to the experimental stimulus *s*(*t*). While this may be a reasonable assumption to make for early sensory processing regions, it may not be true for regions having more complex and nuanced functions such as the default mode networks.

Fig. 2(c), shows that how the observed inter-trial variability in the BOLD signal may arise due to the variability in the HRF. Fig. 2(d) demonstrates that the observed inter-trial variability in the BOLD signal may be due to the variability in the neural activity. In practice, the observed variability may be a combination of both these factors.

If the neural hypothesis is correct, a suitably developed inter-trial synchronization metric may denote the agility of the individual in *disengaging* from the internal mentation or environment monitoring to quickly focus on the task at hand. Such a metric should be correlated to cognitive measures of attention. If found true, this may further lead to the development of fMRI-based measures of mind wandering, concentration or attention.

## Conclusions

We studied the DMN synchronization within trials and across subjects on a stimulus-locked task using a new method called Temporal Synchronization Analysis and found that default networks have low synchronization as compared to task-positive networks. Our study adds to current literature that the DMN should not be considered only as a task-negative network. The inter-trial variations in synchronization may have neural origins as compared to HRF or structural variations which would be more evident in the DMN. General Linear Model based analysis fails to capture such effects and the field should be using other statistical approaches to analyze task fMRI datasets. A method to quantify synchronization can also help measure attentional differences across healthy subjects and even populations with ADHD or Autism.

## Methods

To compare the synchronization across subjects, following publicly available datasets related to different cognitive modalities were used which had the same stimulus onset times across the subjects.

### Dataset

Datasets Used:

- **Food vs Non Food (FNF)**: Photos of food and non-food items were shown to 30 subjects to test craving for food items. Each run had 16 blocks with a 30-second block length and TR of 1.6 seconds [69].
- **Affective Videos (AV)**: Eleven subjects were shown audiovisual stimuli of various emotional valences. Each block had 5 seconds of the stimulus with 7 seconds of fixation. TR was 2.2 seconds [70].
- **Bilingual words (BW)**: Cross-Language repetition priming was tested on 13 bilingual subjects [71] who were shown words of various levels of difficulty in Spanish and English, and they had to rate whether they know it or not. It was a trial based task with 1.5 seconds of stimulus with around 1-6 seconds of fixation and TR of 2 seconds.
- **Human Connectome Project (HCP)**: 169 subjects with four tasks from the HCP Young Adult dataset [72] are used: Social, Emotion, Gambling and Relational. These subjects had both 7T and 3T acquisition and scanned for all seven tasks along with resting-state and movie watching dataset. The four tasks are block-based with 15-25 seconds duration per block and meant to activate various task specific regions.

Study details, participants and MRI acquisition can be referred to individual papers [69; 70; 71; 72] Informed consent was obtained from all subjects. The FNF study was approved by the Medical Ethical Committee of the University Medical Center Utrecht, The Netherlands, the AV study by the Institutional Review Board at the University of South Carolina and HCP study by Washington University institutional review board. The datasets are publicly available on openneuro.org with accession numbers ds000157, ds000205 and ds000051 for FNF, AV and BW respectively and on ConnectomeDB (https://db.humanconnectome.org) for the HCP dataset.

### Preprocessing

We used FSL for preprocessing [73]. The brain was extracted from the high resolution structural file (1mm isotropic) using a repeated version of the FSL-BET. Slice-time correction, motion correction, temporal high-pass filtering (100s), spatial smoothing with 6mm FWHM was done on all datasets. This was followed by co-registration across the dataset and normalization with an MNI 2mm template. For the HCP dataset, the already preprocessed files for task data as available on ConnectomeDB (https://db.humanconnectome.org) which follows the minimal preprocessing pipeline [74] in the CIFTI grayordinates space were used in the analysis.

### GLM analysis

FSL Subject level analysis were done on the subject according to the given task condition. We limited ourselves to first run from the AV dataset and BW dataset and all runs from FNF followed by higher level analysis on the datasets individually. FDR correction (q<0.05) was done to find corrected FDR values. The final maps were the negative logarithm of the corrected FDR values.

We used the FSL Feat tool to analyze the data [73], all the environment variables were taken to create the design matrix and the contrast was active against the fixation. We converted the zstat to p-values which then were FDR corrected using statsmodels toolbox in Python [75]. Negative logarithm was done on these FDR values. The sign of zstat value was multiplied to give us direction information post FDR correction.

For the HCP dataset, we utilized the processed FEAT files as available on ConnectomeDB. For the social task, we limited our analysis to the Theory of Mind (TOM) blocks, face blocks for emotion task, punish blocks for gambling task and match blocks for the relational task.

### Overlap Metrics

The metrics used in table 1 are defined here: (ISC - denotes the number of activated voxels in ISC, TP - the number of task-positive voxels, TN - the number of task-negative voxels)

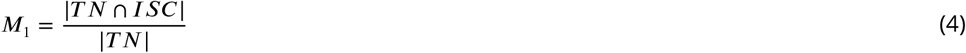

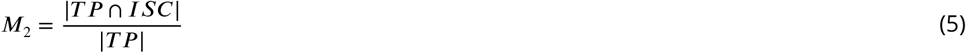

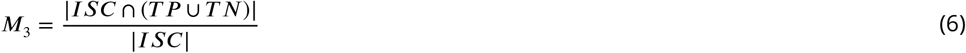

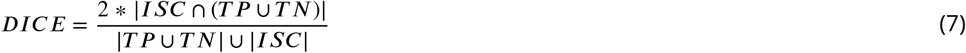

### Temporal Synchronization Analysis (TSA)

The Temporal Synchronization Analysis (TSA) measures the synchronization among a set of signals at different time steps. Let *S*_*i*_(*t*), *i* ∈ [1 … *N*] be a set of *N* signals. The *temporal synchronization* of these signals at time *t* with respect to a baseline *t*_0_, is computed by a two-sided test of hypothesis of percent signal change at time *t* with respect to the baseline signal at time *t*_0_, under the null hypothesis that the percent signal changes are normally distributed with zero mean. Let *Ŝi*(*t*) = (*S*_*i*_(*t*) - *S*_*i*_(*t*_0_))/*S*_*i*_(*t*_0_) * 100. Under the null hypothesis, it is assumed that percent signal change with respect to the baseline, *Ŝi*(*t*) has a normal distribution with zero mean and is independent for all *i* ∈ [1 … *N*]. A two-sided t-test is done, and the corresponding p-value is converted to a standardized Z-statistic *Z*(*t*) = *t*_*to*_*z*(*mean*_*i*_[*Ŝi*(*t*)]/*std*_*i*_[*Ŝi*(*t*)] (where mean and std respectively represent the mean and standard deviations and t_to_z represents the function converting the t-statistic value to the corresponding z-statistic value. The temporal synchronization at time *t* is given by the value of the function *Z*(*t*).

A negative but significant value of temporal synchronization *Z*(*t*) at time *t* indicates that the signals *S*_*i*_(*t*) for *i* ∈ [1 … *N*] synchronously decrease at time *t* with respect to their baseline values at time *t*0. Similarly, a positive but significant value of *Z*(*t*) indicates that all the signals synchronously increase in value with respect to their respective baseline values.

The temporal synchronization may be computed for different trials leading to *inter-trial temporal synchronization (IT-TSA)*, different experimental blocks leading to *inter-block temporal synchronization*, different subjects leading to *inter-subject temporal synchronization (IS-TSA)* or *inter-voxel temporal synchronization (IV-TSA)*.

Let *B*(*e, v, s, t*) represent the BOLD signal of voxel *v*, subject *s*, after a time *t* of the event *e*. The event could be a trial of a particular type or the onset of a specific type of block. The inter-trial temporal synchronization analysis (IT-TSA) for voxel *v* of subject *s* in a set of trials is given as *Z*(*v, s, t*) by computing the temporal synchronization among the signals *B*(*e, v, s, t*) for the events *e* ∈. Similarly, the inter-subject temporal synchronization analysis (IS-TSA) for a set of subjects *U* with respect to a baseline event *e* for a voxel *v*, is given as *Z*(*e, v, t*) by computing the temporal synchronization among the signals *B*(*e, s, t, v*) for the subjects *s* ∈ *U*. In a similar vein, the inter-voxel temporal synchronization analysis (IV-TSA) for a set of voxels *V*, with respect to a baseline event *e*, for a given subject *s*, is given as *Z*(*e, s, t*) by computing the temporal synchronization among the signals *B*(*e, v, s, t*) for the voxels *v* ∈ *V*.

In practice, the inter-trial temporal synchronization analysis (IT-TSA) is expected to be significant only for the voxels that get activated (or deactivated) due to the event and is expected to become insignificant (unless the subsequent trials or blocks start interfering) as soon as changes in the expected BOLD response becomes close to zero (around 15-20 seconds after the end of the block or trial). The TSA offers a viable alternative to the conventional General Linear Model (GLM) analysis of the fMRI data. Unlike GLM, TSA does not make the linear time invariance (LTI) assumption, nor does it assume any pre-defined shape of hemodynamic response function (HRF). Instead, TSA is completely model-free and may be used to discover double-peak behaviour [76] as well as shape of the hemodynamic response function HRF [47] for different voxels. However, given that IT-TSA doesn*’*t combine different time points after the event or block, it may need a larger number of trials as compared to conventional GLM analysis to achieve similar levels of significance.

For naturalistic paradigms such as those used in Neurocinematics [27; 28] or other similar experiments [29; 37], the TSA offers a much better alternative to the Inter-Subject Correlation analysis generally used for the analysis of the fMRI data under such paradigms. In such paradigms, the stimulus presented to different subjects is perfectly aligned and the inter-subject correlations (ISC) are computed to infer brain activations. The inter-subject temporal synchronization analysis (IS-TSA) gives an instantaneous view of synchronized activations in different voxels across multiple subjects, whereas the ISC analysis only gives the aggregate activations for the duration of the experiment. Thus, using the IS-TSA it is possible to not only find which voxels get activated, but also at what time instants during the experiments they get activated. Moreover, even in the conventional experimental paradigms, when the stimulus presented to different subjects are not perfectly aligned, it is possible to use IS-TSA in a manner similar to IT-TSA, by using a subject-specific baseline that is aligned with the subject-specific stimulus or event presentations. In this case, multiple repetitions of the stimulus are not needed to get robust statistics. The IS-TSA can be used to find the entire course of activations of different voxels even for a single event.

The IV-TSA provides an alternate method to carry out analysis similar to the regional homogeneity (REHO) [77] for fMRI data. For every voxel, a block of voxels in its neighbourhood maybe used to compute the inter-voxel temporal synchronization analysis (IV-TSA). For this analysis, a baseline such as the start of a block or event or average BOLD signal may be used. Thus, for every voxel, IV-TSA may be computed among its neighbouring voxels (say within a distance of 15mm or in a 3×3 neighbourhood box), at every time point using the baseline. Thus, IV-TSA gives a temporal view of the instantaneous synchronization among the neighbouring voxels which is expected to be closely related to the voxel activations during the presentation of the events.

The temporal synchronization analysis TSA provides a powerful and unifying framework for statistical analysis of fMRI data which will enable fMRI researchers to make many discoveries. The IT-TSA and IS-TSA, which does not depend on any of the restrictive GLM assumptions, may be used to re-analyze the existing fMRI data to make newer discoveries. The IS-TSA may be used to augment the analysis of fMRI data collected under naturalistic paradigms and provide precise information about temporal sequence of activations of different voxels. It may also be used to re-analyze the fMRI data collected using conventional task-based paradigms, especially for the experiments having a large number of subjects. The IV-TSA may be used to study the temporal dynamics of regionally synchronized BOLD activity at a temporal resolution (as less as 1 TR) that was not possible earlier.

### T-test analysis

Independent samples t-test analysis was run on the IT-TSA using Scipy toolbox [78] values to determine differences across the synchronization using different ROIs. The max IT-TSA values across time was averaged across subjects and then compared (t-test) across the voxels in the task-positive and default mode network ROIs from the Schaefer atlas which are depicted in Fig. 6.

### Linear Regression

We used the statsmodels python package [75] to fit Ordinary Least Squares (OLS) regression models to the scatter plots.

### Correlation analysis

To compute the efficacy of the IS-TSA metric, we computed the IS-TSA per voxel and then correlated with the HRF convolved stimulus and averaged the Pearson correlation across the ROI and shown in Fig. 10.

### Intersubject correlation (ISC)

ISC analysis was done with scripts written in Python [79]. Voxel wise correlations across subjects are averaged, we get a final ISC map with correlation values. Correlations were computed as given below using method provided in [27; 35]

#### Correlation computation

To perform ISC, we calculate the correlations in the time series of a single voxel across all pairs of subjects.

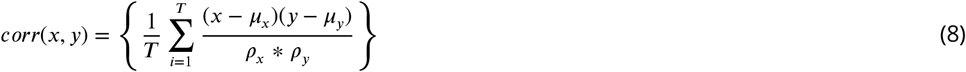

Where:

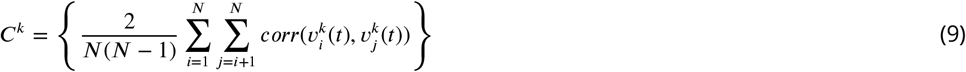

Where:

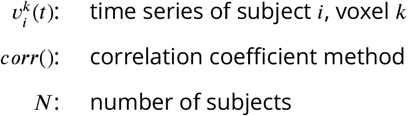

## Data/Code Availability Statement

All datasets are publicly available from https://openneuro.org and https://db.humanconnectome.org and codes are available at bitbucket repository https://bitbucket.org/vtripathi/iisc/src/master/

## Acknowledgements

The authors would like to thanks Jaspreet Kaur and Varun Kumar for their valuable comments and inputs. Box figures were created using Biorender. We would like to acknowledge the OpenfMRI project and NSF Grant OCI-1131441 for the publicly available datasets upon which we conducted the analyses.

## Author contributions statement

V.T conducted the experiments. V.T. and R.G. analyzed the results and worked on the manuscript.

